# Depletion of border-associated macrophages impairs CSF flow and exacerbates extranuclear mutant Huntingtin aggregation in the zQ175DN mouse model of Huntington’s disease

**DOI:** 10.64898/2026.09.24.754172

**Authors:** Yanli Jiang, Rahul Sondawle, Helen Moniz, Emily Mu, Wenzhen Duan

## Abstract

Huntington’s disease (HD) is a neurodegenerative disorder driven by a CAG trinucleotide repeat expansion in the *huntingtin* (*HTT*) gene, resulting in neuronal degeneration. While cell-autonomous mechanisms within neurons have been extensively characterized, how brain waste clearance pathways at the brain’s borders change during disease progression remains poorly understood. In this study, we investigated the structural and functional adaptations of border-associated macrophages (BAMs) in the zQ175DN knock-in mouse model of HD. Immunohistochemical analysis revealed an age-dependent, selective reduction in the CD206⁺ Lyve1⁺ subset of leptomeningeal BAMs in 16-month-old zQ175DN mice, which tightly correlated with a profound impairment in cerebrospinal fluid (CSF) flow velocity and perivascular influx. To determine whether BAMs actively drive this CSF flow, we selectively depleted these populations of macrophages in 10-month-old zQ175DN mice via intracerebroventricular (i.c.v.) injection of clodronate liposomes (CLO). Chemical ablation of BAMs significantly exacerbated global CSF dynamic impairment, indicted by decreasing CSF fluorescent tracer distribution across perivascular parenchymal channels. Crucially, this CSF flow failure resulted in a marked accumulation of extranuclear mutant huntingtin (mHTT) aggregates in the zQ175DN HD mouse brain, while intranuclear inclusions were not significantly altered. Furthermore, Acute BAMs depletion had no impact on microglial or astrocytic activation in the brain, demonstrating that the exacerbation of mHTT load is a byproduct of brain waste clearance dysfunction rather than gliosis. Together, our results indicate that BAMs are non-redundant regulators of CSF flow dynamics at the brain borders, offering a unique non-neuronal mechanism to mHTT accumulation and potential a therapeutic target to combat HD.

## Introduction

Huntington’s disease (HD) is a progressive, inherited neurodegenerative disorder clinically characterized by chorea, cognitive decline, and psychiatric disturbances, pathologically highlighted by severe atrophy of the striatum and cerebral cortex. The molecular etiology centers on a CAG trinucleotide repeat expansion within exon 1 of the *huntingtin* (*HTT*) gene, yielding a mutant huntingtin (mHTT) protein containing an expanded polyglutamine (polyQ) tract [1]. While intracellular cascades, such as transcriptional dysregulation, mitochondrial failure, and synaptic vesicle transport disruption, have been extensively investigated within neurons, therapeutic strategies targeting these cell-autonomous mechanisms have yielded limited clinical success by far [2, 3]. Consequently, attention has expanded toward systemic structural vulnerabilities, particularly recently recognized brain waste clearance systems which is functioning to eliminate toxic proteins and metabolic wastes from the central nervous system.

In physiological states, metabolic debris and toxic proteins are continuously cleared from the parenchyma through a coordinated interstitial fluid (ISF) network, described as glymphatic system [4, 5]. This network relies on the convective influx of cerebrospinal fluid (CSF) along periarterial spaces, its mixing with ISF, and subsequent efflux along perivenous pathways and meningeal lymphatic channels to the peripheral lymph nodes [6, 7]. Dysfunction within this fluid exchange system is a hallmark of several neurodegenerative proteinopathies, including Alzheimer’s and Parkinson’s diseases, where impaired fluid dynamics accelerate the extracellular seeding and aggregation of toxic protein species [7, 8]. Emerging evidence suggests that glymphatic clearance rates are similarly suppressed in mouse models and clinical carriers of HD, yet the underlying cellular drivers responsible for this clearance system failure remain fully understood [9–11].

A recent study reported that prominent candidates regulating the CSF flow via glymphatic channels are border-associated macrophages (BAMs) [12]. Unlike parenchymal microglia, BAMs are long-lived, self-renewing myeloid cells residing at strategic fluid-brain interfaces, including the leptomeninges, perivascular spaces, and choroid plexus [13, 14]. BAMs are phenotypically distinguished by their stable expression of the mannose receptor CD206 [15]. They are sub-categorized based on localized expression signatures, such as Lyve1 (lymphatic vessel endothelial hyaluronan receptor 1), which marks subsets aligned with lymphatic drainage pathways [16]. Functionally, BAMs act as metabolic gatekeepers; they clear cellular debris, monitor vascular permeability, and regulate vascular tone through extracellular matrix remodeling [17, 18] . Despite their critical positioning at the main checkpoints of fluid entry and exit, how BAM networks adapt over the lifespan in HD, and whether their dysfunction directly compromises CSF mechanics and mHTT aggregate loading, remains entirely unknown.

To address these questions, we leveraged the zQ175DN knock-in mouse model, which recapitulates the progressive onset of HD pathology including mHTT aggregates accumulation [19]. We mapped the phenotypic distribution of BAMs subpopulations across pre-symptomatic (3 months), onset of symptomatic (6 months), and advanced disease stages (16 months) in this HD mouse model. We identified an age-dependent decrease in parenchymal BAMs and Lyve1-positive BAMs in the leptomeninges of zQ175DN HD mice. The alterations of BAMs accompanied by compromised CSF flow dynamics. Chemical depletion of BAMs in adult HD mice further exacerbates CSF flow impairment, suggesting a causal link between BAMs and CSF flow regulation in HD condition. Furthermore, BAMs depletion exacerbates extranuclear mHTT aggregate load, while this manipulation had no influence on gliosis. Our findings demonstrate that losing BAM negatively regulates global and parenchymal CSF flow dynamics, accelerating the accumulation of toxic mHTT aggregates independently on neuroinflammatory responses.

## Results

### Age-dependent reduction of leptomeningeal and parenchymal border-associated macrophages (BAMs) in HD mice

To evaluate the spatial distribution and cellular architecture of BAMs networks in the HD, we performed immunohistochemistry with BAMs specific markers in zQ175DN mice and their littermate wild type (WT) controls. Co-labeling for the vascular basement membrane component Collagen IV and the pan-BAMs marker CD206 verified that BAMs are positioned within the structural architecture of the leptomeninges and embedded within the adventitial layers of superficial and penetrating cerebral blood vessels in HD and WT mice (**Supplementary Fig. 1A, B**).

To track alterations across the disease trajectory, we quantified total BAMs counts across three ages: 3 months (pre-symptomatic), 6 months (early symptomatic), and 16 months (advanced disease state). Broad spatial quantification of total CD206⁺ cells tracking along the leptomeningeal borders showed stable cell numbers across all time points, with no statistically significant differences observed between WT and zQ175DN mice (**Fig. 1A, B**). Conversely, quantification of deep parenchymal, perivascular CD206⁺ populations revealed a clear, age-associated decline across the mouse lifespan in both genotypes (**Fig. 1C**). This loss of BAMs population was driven primarily by natural aging rather than the *HTT* mutation itself, as shown by significant decreases at 16 months relative to the 3-month baseline in both WT (P < 0.01) and zQ175DN lines (P < 0.05), with no statistical differences seen directly between genotypes within identical age brackets (**Fig. 1C**).

**Figure 1.**
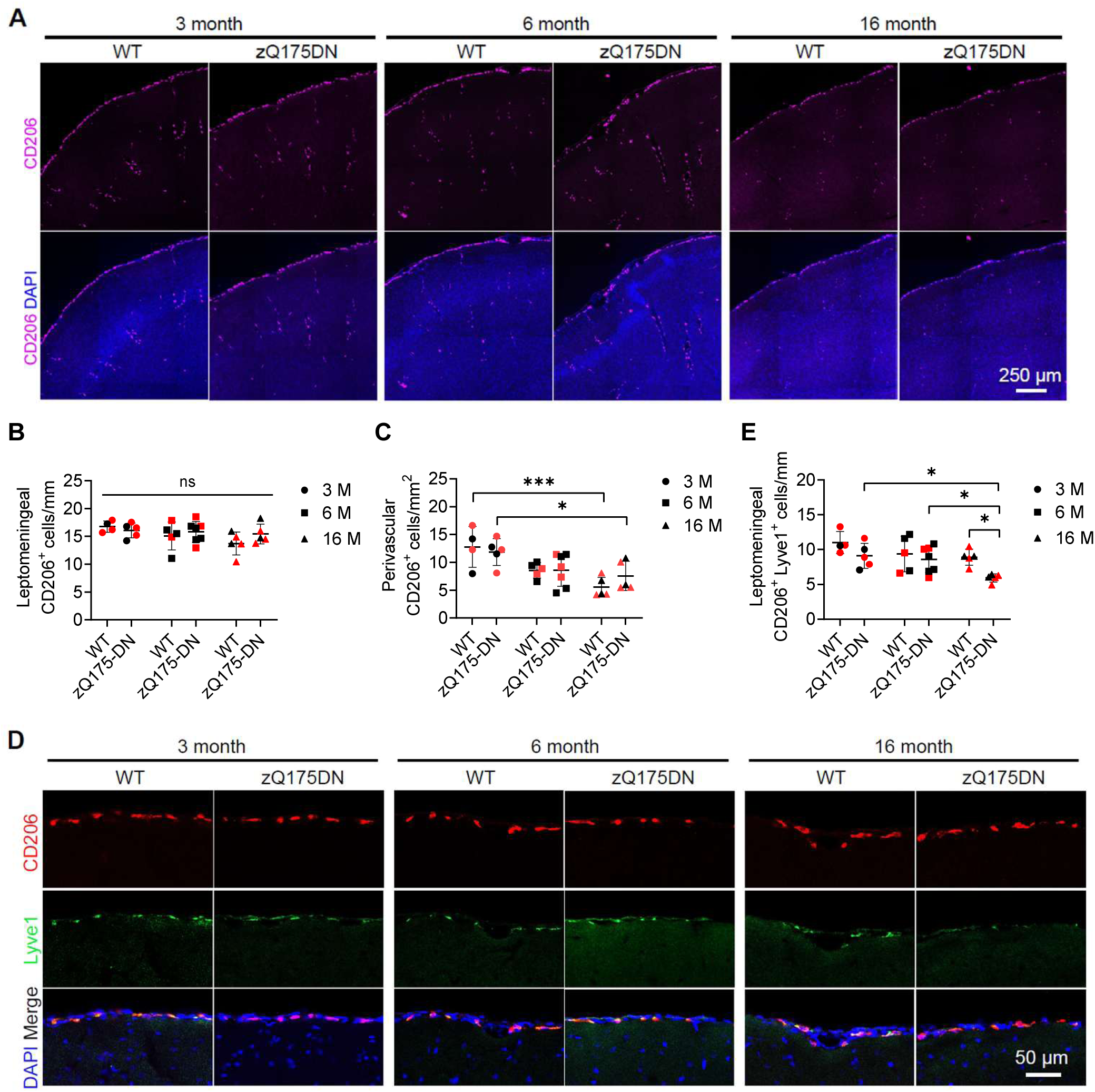
Leptomeningeal Lyve1-positive border associated macrophages (BAMs) are selectively decreased in aged HD mice. **(A)** Representative confocal images showing CD206⁺ border-associated macrophages (magenta) counterstained with DAPI (blue) in the brains of WT and zQ175DN mice at 3, 6, and 16 months of age. Scale bar = 250 μm. **(B)** Quantitative analysis of absolute leptomeningeal BAMs linear density (CD206⁺ cells/mm) along the outer meningeal borders across the indicated age cohorts. Ordinary two-way ANOVA revealed no significant main effect of genotype, F (1, 25) = 0.7555, P = 0.3930, no significant genotype × age interaction, F (2, 25) = 0.9996, P = 0.3823; and no significant main effect of age, F (2, 25) = 2.272, p = 0.1240. Therefore, no post hoc multiple comparisons were performed. **(C)** Quantitative analysis of deep parenchymal perivascular BAMs numbers (CD206⁺ cells/mm²) across indicate ages. Data were analyzed using ordinary two-way ANOVA. There was no significant main effect of genotype, F(1, 25) = 0.1732, P = 0.6808, or genotype × age interaction, F (2, 25) = 0.8198, p = 0.4520. However, there was a significant main effect of age, F (2, 25) = 12.23, P = 0.0002. Therefore, Sidak’s post hoc multiple comparisons test was performed; ns = not significant, *P < 0.05, **P < 0.01. **(D)** Representative CD206 (red) and Lyve1 (green) co-immunostained confocal images showing specialized leptomeningeal Lyve1⁺ BAMs networks counterstained with DAPI (blue) in WT and zQ175DN brains across the lifespan. Scale bar = 50 μm. **(E)** Quantitative analysis of leptomeningeal CD206⁺ Lyve1⁺ double-positive cell numbers per millimeter (cells/mm) demonstrating a selective reduction at 16 months of age in HD mice. Data were analyzed using ordinary two-way ANOVA. There were significant main effects of genotype, F (1, 25) = 8.973, P = 0.0061, and age, F (2, 25) = 5.599, P = 0.0098. However, the genotype × age interaction was not significant, F (2, 25) = 1.083, P = 0.3539. Tukey’s post hoc multiple comparisons test was performed to identify difference between age groups. *P < 0.05. For all graphs, individual data points are color-coded by sex: red symbols indicate female mice, black symbols indicate male mice; 3 months: n = 4 WT (3F/1M), n = 5 zQ175DN (3F/2M); 6 months: n = 5 WT (2F/3M), n = 7 zQ175DN (3F/4M); 16 months: n = 5 WT (3F/2M), n = 5 zQ175DN (3F/2M). Data presented as mean ± SD.

We next examined the specific subpopulation of leptomeningeal BAMs double-positive for Lyve1. While these subtype BAMs remained stable at 3 and 6 months of age, we noted a statistically significant, selective reduction (P < 0.05) in CD206⁺ Lyve1⁺ cell numbers exclusively in 16-month-old zQ175DN mice relative to age-matched WT controls (**Fig. 1D, E**).

### Impaired cerebrospinal fluid (CSF) dynamics in aged HD mice

Given that the age-related depletion of the CD206⁺ Lyve1⁺ subpopulation occurred at an advanced disease stage of zQ175DN HD mice (16 months), we tested whether CSF transport was altered in parallel. We performed intracerebroventricular (i.c.v.) microinjections of a fluorescent CSF flow tracer, BSA-647, into 16-month-old WT and zQ175DN mice and tracked CSF flow dynamics over a 60-minute time-lapse using *in vivo* stereoscopy live imaging (**Fig. 2A**).

**Figure 2.**
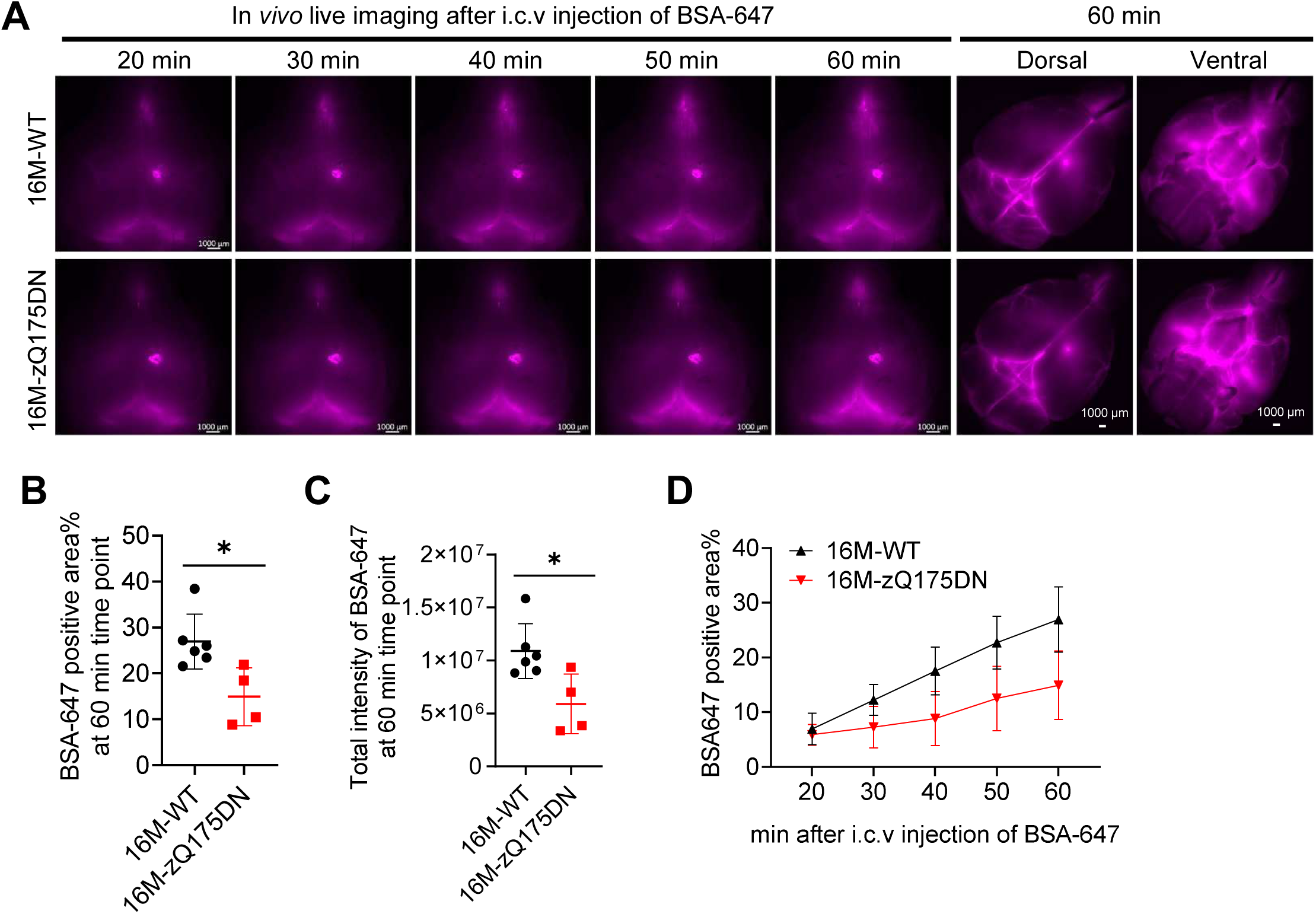
Impaired CSF tracer influx into the brain parenchyma in aged HD mice. **(A)** Representative time-lapse fluorescence macro-images tracking the surface distribution and clearance velocity of i.c.v.-injected far-red fluorescent tracer (BSA-647; magenta) across the cranium of 16-month-old WT and zQ175DN mice at 20, 30, 40, 50, and 60 minutes post-injection. Isolated ex vivo dorsal and ventral macro-views at the 60-minute endpoint are provided on the right. Scale bars = 1000 μm. **(B-C)** Quantification of the total percentage of BSA-647-positive area (B) and absolute integrated fluorescence intensity (C) at the 60-minute endpoint. (**D**) Real-time kinetic curve tracking the percentage of BSA-647-positive surface area over the entire 60-minute observation window is shown on the right. Data are from n = 6 WT mice (4 females, 2 males) and n = 4 zQ175DN mice (2 females, 2 males). Data are presented as individual points or mean ± SD. Statistical comparisons between genotypes were evaluated using an unpaired two-tailed Student’s t-test at each respective time point; *P < 0.05.

Fluorescence mapping revealed a stark constraint in tracer migration across the calvaria surface in the HD line compared to WT controls (**Fig. 2A**). Quantitative tracking across the 60-minute trial demonstrated that both the percentage of BSA-647-positive area and the total integrated fluorescence intensity were significantly reduced in 16-month-old zQ175DN brains compared to WT controls at the 60-minute mark (P < 0.05; **Fig. 2B, C**). Real-time kinetic analysis further validated this CSF flow disturbance, displaying a consistently lower tracer-positive area surface profile in the zQ175DN cohort throughout the entire 60-minute imaging period (P < 0.05; **Fig. 2D**).

### BAMs depletion exacerbates CSF flow failure in HD mice

Because the loss of specialized leptomeningeal BAMs occurred alongside a decline in CSF tracer flow, we hypothesized that BAMs are actively required to maintain CSF flow efficiently. To establish a direct causal link, we chemically depleted BAMs populations by injecting clodronate liposomes (CLO) via an i.c.v. route into 10-month-old zQ175DN mice, choosing an age preceding natural BAMs decline (**Fig. 3A**). Mice were sacrificed two weeks post-injection to verify BAMs ablation. Confocal imaging of meningeal mounts immunolabeled for CD206 demonstrated an almost complete destruction of the leptomeningeal BAMs in clodronate-treated mice (**Fig. 3B**). Quantification of remaining cells confirmed a highly significant, comprehensive reduction in leptomeningeal BAM density in the zQ175DN mice recevied CLO compared to those zQ175DN mice received vehicle (P < 0.0001), validating the efficacy of CLO-induced depletion of BAMs in HD mice (**Fig. 3C**).

**Figure 3.**
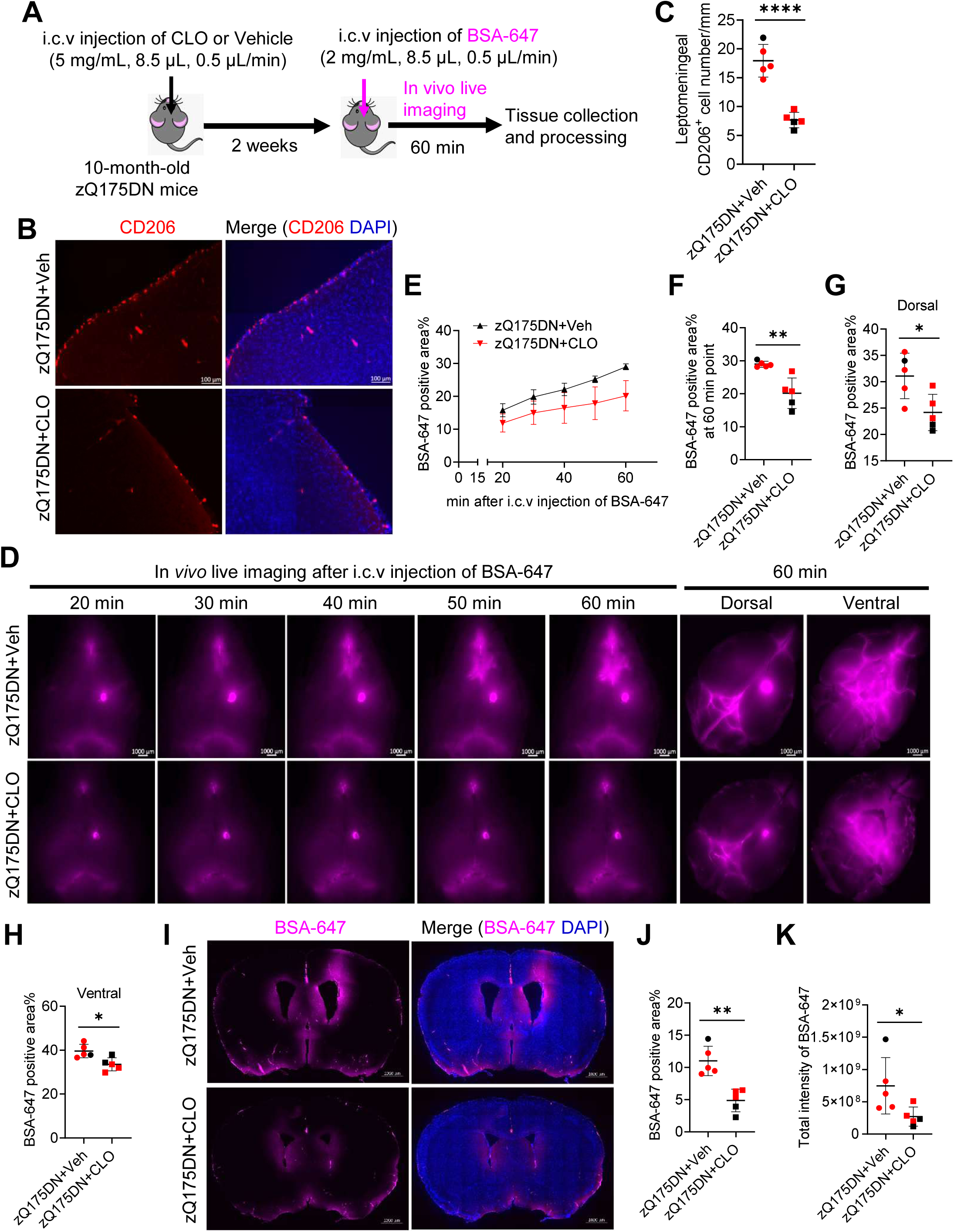
Border-associated macrophages (BAMs) depletion directly impairs parenchymal CSF flow dynamics in HD mice. **(A)** Detailed experimental protocol timeline: 10-month-old zQ175DN mice received an i.c.v. injection of clodronate-loaded liposomes (CLO) or vehicle control; 2 weeks later, functional clearance was measured via i.c.v. injection of BSA-647 over a 60-minute imaging period prior to tissue collection. **(B)** Representative CD206 immunofluorescence staining (red) counterstained with DAPI (blue) in zQ175DN cortical borders 2 weeks post-injection, validating chemical network ablation. Scale bar = 100 μm. **(C)** Statistical quantification of total remaining leptomeningeal CD206⁺ cells per millimeter verifying comprehensive knockdown efficiency. Unpaired two-tailed t-test; ****P < 0.0001. **(D)** Representative time-lapse images showing macroscopic cranial surface delivery of BSA-647 tracer (magenta) over 60 minutes, with isolated 60-minute dorsal and ventral whole-brain views shown on the right. **(E)** Kinetic tracking curves charting the cumulative percentage of BSA-647-positive area across the 60-minute window. **(F-H)** Quantitative analysis of the percentage of tracer-positive surface area at the 60-minute session endpoint across the entire whole brain (F), isolated dorsal view (G), and isolated ventral view (H). Unpaired two-tailed t-test; *P < 0.05, **P < 0.01. **(I)** Representative ex vivo coronal brain slice maps tracking deep parenchymal and perivascular tracer penetration (magenta) counterstained with DAPI (blue) at 60 minutes post-injection. Scale bar = 1000 μm. **(J-K)** Quantitative tracking of the percentage of tracer-occupied sectional area (J) and integrated total fluorescence tracer intensity (K) within internal cerebral sections. Unpaired two-tailed t-test; *P < 0.05, **P < 0.01. For all graphs, red symbols indicate female mice, black symbols indicate male mice; vehicle control group: n = 5 (3F/2M), CLO treatment group: n = 5 (3F/2M). Data presented as mean ± SD.

Following the two-week chemical ablation period, we evaluated CSF transport mechanics by tracking i.c.v.-infused BSA-647 tracer dynamics over a 60-minute period (**Fig. 3D**). Real-time analysis of CSF flow influx demonstrated that zQ175DN mice lacking BAMs networks (CLO cohort) exhibited a severe restriction in tracer dispersion across the glymphatic channels compared to vehicle-treated group (**Fig. 3E**). Quantitative endpoint analysis at 60 minutes post-injection confirmed a significant decrease in global tracer distribution area (P < 0.01; **Fig. 3F**). This reduction was driven by substantial CSF constraints across both the dorsal (P < 0.05; **Fig. 3G**) and ventral (P < 0.05; **Fig. 3H**) cerebrum.

To examine CSF flow into deep parenchyma, we evaluated coronal brain slices harvested at the 60-minute post tracer injection (**Fig. 3I**). Brain sections from the zQ175DN mice received CLO displayed significantly decreased tracer penetration along perivascular channels flanking the lateral ventricles and leptomeninges (**Fig. 3I**). Quantification confirmed a significant reduced distribution of tracer, indicated by tracer-positive sectional area (P < 0.01; **Fig. 3J**) and the corresponding total fluorescence intensity (P < 0.05; **Fig. 3K**) within parenchyma, indicating that removing BAMs directly worsens CSF flow dynamics in the HD brain.

### BAMs loss selectively increases extranuclear mHTT accumulation

We next evaluated whether BAMs depletion and associated CSF flow impairment impacted mHTT clearance and altered toxic aggregate load. Brain sections were immunolabeled using EM48 antibody to label mHTT inclusions (**Fig. 4A**) (DiFiglia et al., 1997). Confocal imaging analysis revealed prominent, dense nuclear mHTT aggregates distributed throughout the striatum (**Fig. 4A**). Stripping the BAMs via clodronate liposomes in 10 months old zQ175DN mice had no significant influence on the nuclear mHTT inclusions (**Fig. 4B**). In contrast, the density of neuropil-localized, extranuclear mHTT aggregates were significantly increased in zQ175DN mice received CLO compared to HD mice with vehicle controls (P < 0.05; **Fig. 4C**) (Aronin and Sapp, 2024). This selective accumulation indicates that BAMs-driven fluid currents are actively required to clear extranuclear mHTT from the brain parenchyma in the stage when disease is already started.

**Figure 4.**
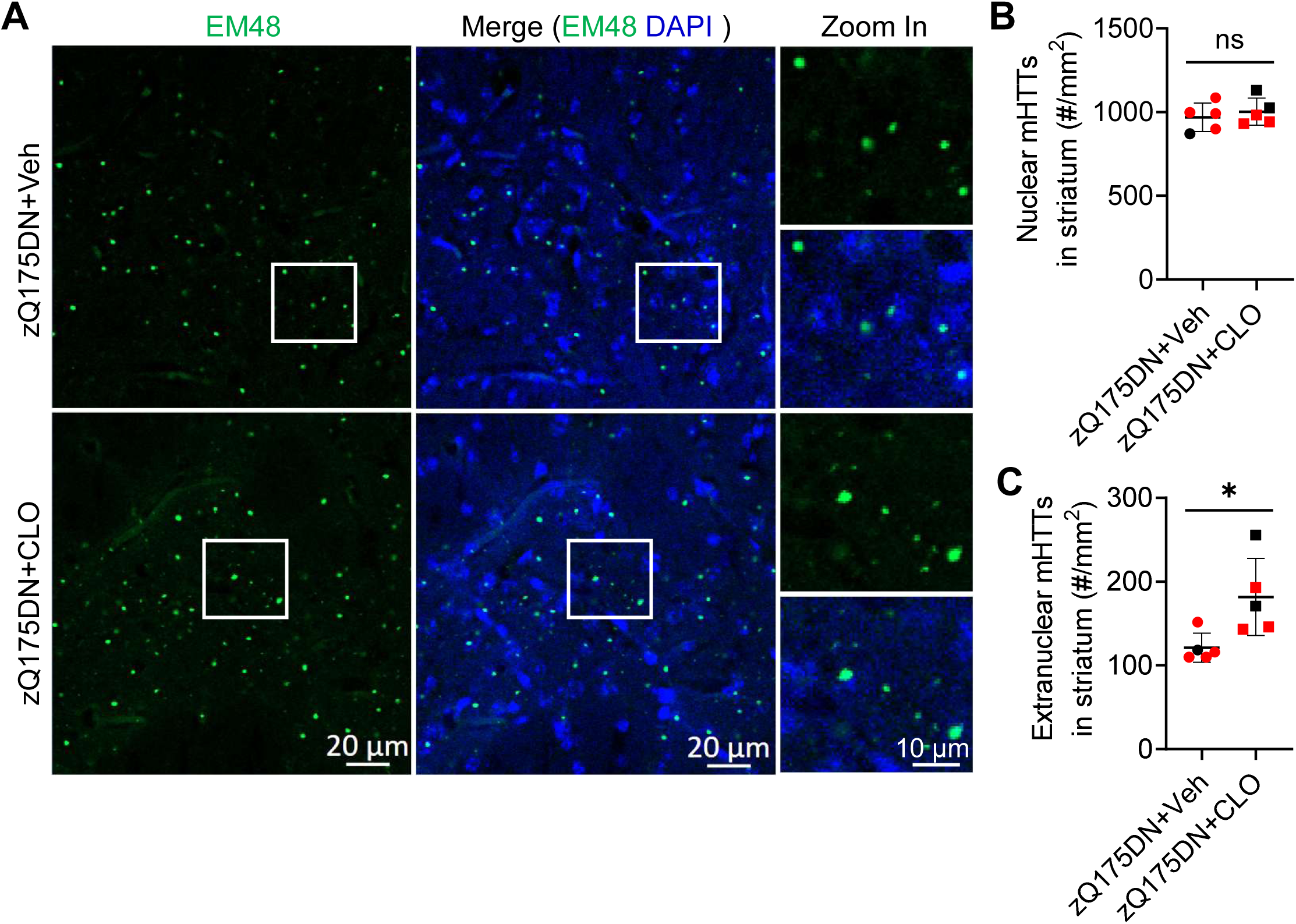
Border-associated macrophages (BAMs) depletion significantly exacerbates extranuclear mHTT aggregate load in HD mice. **(A)** Representative confocal images showing EM48-stained mutant huntingtin (mHTT) aggregates (green) counterstained with DAPI (blue) within the striatal matrix of 10-month-old zQ175DN mice 2 weeks after i.c.v. vehicle or CLO delivery. White framed regions are magnified on the right to track isolated aggregates in the neuropil. Scale bars = 20 μm (main panels) and 10 μm (high-magnification zoom). **(B)** Statistical quantification of structural nuclear mHTT aggregate density (number of inclusions per square millimeter; #/mm²) showing no significant alterations following treatment. **(C)** Quantification of neuropil-localized, extranuclear mHTT aggregate density (#/mm²) demonstrating an isolated increase after BAMs ablation. Red symbols represent female mice, black symbols represent male mice; vehicle group: n = 5 (3F/2M), CLO group: n = 5 (3F/2M). Data analyzed via an unpaired two-tailed Student’s t-test; ns = not significant, *P < 0.05. Data presented as mean ± SD.

### BAMs depletion does not alter astrocyte or microglial gliosis in HD mice

Finally, to rule out whether the increased mHTT load and impaired CSF flow were caused by generalized neuroinflammation, we evaluated classical neuroinflammatory cellular metrics. Striatal and cortical areas were labeled for GFAP (astrocytes) and Iba1 (microglia) (**Fig. 5A**). Morphological profiling showed no obvious differences in glial distribution or activation state between clodronate-treated and vehicle-treated HD mice (**Fig. 5A**). Detailed statistical analysis confirmed that removing BAMs did not cause secondary microgliosis in both brain regions we examined (**Fig. 5B-E**). There was no indication of astrogliosis caused by BAMs depletion, indicated by no changes in percentage of total GFAP positive area (**Fig. 5F**), integrated total fluorescence intensity (**Fig. 5G**) for GFAP, number of GFAP-positive astrocytes within both the cortex (**Fig. 5H**) and striatum (**Fig. 5I**). These data indicate that the mHTT accumulation observed after BAMs ablation is a consequence of impaired CSF clearance pathway, rather than a byproduct of altered gliosis.

**Figure 5.**
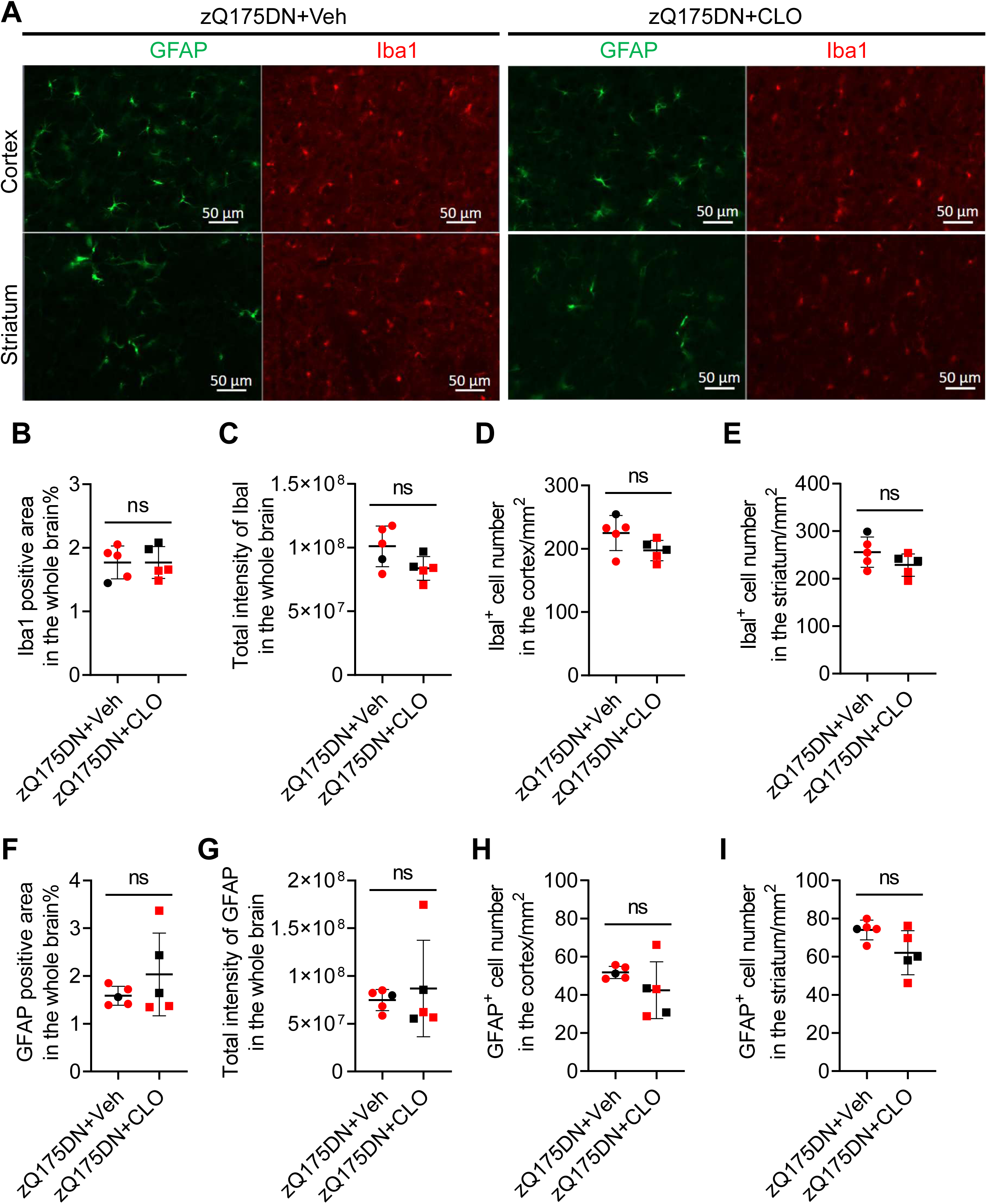
Depletion of border-associated macrophages (BAMs) does not induce reactive microglia and astrocyte gliosis in HD mice. **(A)** Representative immunofluorescence confocal tile scans showing GFAP⁺ activated astrocytes (green) and Iba1⁺ reactive microglia (red) within the cerebral cortex and striatal matrix of zQ175DN mice 2 weeks after vehicle or CLO injection. Scale bar = 50 μm. **(B, C)** Q Quantitative assessment of the percentage of total Iba1-positive area (B) and integrated total Iba1 fluorescence intensity (C) across the whole brain section. **(D, E)** counting of absolute Iba1-positive cell numbers per square millimeter (#/mm²) within the cerebral cortex (D) and striatal matrix (E). **(F, G)** Quantitative assessment of the percentage of total GFAP-positive area (F) and integrated total GFAP fluorescence intensity (G) across the whole brain section. **(H, I)** Statistical counting of absolute GFAP-positive cell numbers (#/mm²) within the cerebral cortex (H) and striatum (I). Individual data points are plotted: red symbols indicate female mice, black symbols indicate male mice; vehicle cohort: n = 5 (3F/2M), CLO cohort: n = 5 (3F/2M). Data evaluated using an unpaired two-tailed Student’s t-test; ns = not significant. Data presented as mean ± SD.

## Discussion

This study provides direct evidence that BAMs are non-redundant cellular regulators supporting CSF flow dynamics and mHTT aggregate clearance in HD. Using the zQ175DN knock-in mouse model, we demonstrated an age-dependent, selective reduction in the CD206⁺ Lyve1⁺ subpopulation of leptomeningeal BAMs during advanced stages of the disease. This subpopulation of BAM loss matched a decline in CSF perivascular flow and its associated waste clearance function. By using clodronate liposomes to ablate the BAM network prior to their natural aging-related decline, we proved a direct causal link: losing BAMs significantly worsens perivascular CSF inflow [12, 20]. Crucially, this fluid transport failure selectively caused extranuclear, neuropil-localized mHTT aggregates to accumulate in the HD brain, while leaving parenchymal neuroinflammatory responses unaltered.

The selective vulnerability of the CD206⁺ Lyve1⁺ BAM subpopulation in advanced HD highlights a critical cellular shift at the brain borders. Lyve1 expression typically marks BAMs that line the leptomeningeal vasculature and major dural sinuses, positioning them as primary regulators of waste egress into peripheral lymphatic networks [21]. The targeted loss of this population in HD mice at advanced disease stage suggests that chronic exposure to mHTT selectively damaged these border macrophage subsets. Interestingly, our tracking of leptomeningeal BAMs revealed a significant decline of CD206 Lyve1-positive BAMs in HD, suggesting that the leptomeningeal-CSF boundary is uniquely vulnerable to HD-specific pathogenesis, directly compromising CSF transport.

Our BAM depletion experiments clarify the role BAMs play in driving CSF transport in HD. The profound stagnation of the BSA-647 tracer across both external cortical surfaces and within deep perivascular channels in BAMs-depleted mice indicates these cells are required to maintain CSF flow and its associated clearance pathways [12, 22]. BAMs are known to continuously clear extracellular matrix debris, modify basement membranes, and modulate vascular tone through chemical messaging with endothelial cells and vascular smooth muscle cells [13, 23]. Removing this homeostatic support likely causes perivascular channels and meningeal drainage zones to be impaired, raising resistance along ISF-CSF pathways and reducing convective fluid movement [6]. This structural restriction was indicated in our coronal sections, which showed that losing BAMs severely limits tracer delivery into deep parenchymal spaces. This demonstrates that BAMs are actively required to maintain the physical patency of fluid inflow routes into deep brain matrices.

Finally, a major finding of our study is that this increased mHTT aggregate load and CSF flow failure occurred independently of changes in neuroinflammation in HD following BAM depletion. This decoupling demonstrates that the worsening of mHTT pathology by depletion of BAMs is driven by a brain clearance failure rather than gliosis, differentiating CSF fluid dynamics from cellular immune networks [24]. In summary, our results indicate that BAMs are vital cellular pillars required to maintain effective cerebral waste-clearance pathways and protect the brain from toxic protein buildup in HD. Therapeutic strategies aimed at supporting BAMs networks or boosting their functional capacities at CSF flow control could offer a valuable approach to combat toxic protein load in HD.

## Materials and Methods

### Animals

All experimental procedures involving mice were approved by the Johns Hopkins University Animal Care and Use Committee and conducted in accordance with the National Institutes of Health guidelines for the care and use of laboratory animals. For baseline BAMs mapping and CSF flow analysis, male and female WT and zQ175DN mice were evaluated at 3 months (WT: 3 female, 1 male; zQ175DN: 3 female, 2 male), 6 months (WT: 2 female, 3 male; zQ175DN: 3 female, 4 male), and 16 months of age (WT: 3 female, 2 male; zQ175DN: 3 female, 2 male). The zQ175DN KI mice used in this study were backcrossed onto a C57BL/6J background strain (JAX stock #000664). All mice were housed in a specific pathogen-free facility under controlled conditions (23 °C, 30–70% humidity) on a 12 h reversed light–dark cycle, with ad libitum access to food and water. For chemical depletion studies, 10-month-old zQ175DN mice were randomly assigned to either the Vehicle group (4 female, 1 male) or the Clodronate Liposome (CLO) group (3 female, 2 male). Tail biopsies were collected from all mice for genotyping and CAG repeat length determination. Samples were analyzed by Transnetyx Inc.

### BAMs depletion by intracerebroventricular (i.c.v) injection

To deplete BAMs, zQ175DN mice received an intracerebroventricular (i.c.v.) injection of clodronate-loaded liposomes (CLO; 8.5 μL; 5 mg/ml; at a rate of 0.45 μL/min; CLD-8901; Thermo Fisher Scientific) [25] at 10 months of age. The control zQ175DN group received i.c.v. injection of PBS-loaded liposomes (vehicle) at the same age.

For the i.c.v. injection procedure, mice were anesthetized with ketamine/xylazine, and the hair on the head was shaved using a trimmer. Each mouse was then placed on a stereotaxic frame and secured with ear bars on both sides of the head and a nose bar to ensure proper leveling. Once the head was stabilized, the surgical area was sterilized with alternating applications of alcohol and iodine. To prevent corneal drying during the procedure, an ophthalmic ointment was applied to both eyes. Under a dissection microscope, the scalp was lifted with forceps and removed using fine scissors to expose the skull. The Hamilton syringe was positioned at the coordinates Anterior/Posterior (AP) −0.6 mm and Medial/Lateral (ML) −1.2 mm relative to Bregma. The target site was marked by scoring the skull surface with fine forceps. A small hole was drilled at the marked location using a microdrill. The Hamilton syringe was then carefully lowered to the brain surface to establish the dorsal–ventral (DV) reference point and then advanced to DV −2.0 mm. Using a syringe pump, 8.5 μL of CLO or vehicle was infused at a rate of 0.45 μL/min. Following injection, the syringe was left in place for 10 minutes to facilitate diffusion and was then slowly withdrawn. The scalp was sutured and disinfected, after which the mouse was placed on a heating pad until full recovery and subsequently returned to its home cage. To evaluate the efficiency of CLO–liposomes, BAMs cell numbers were quantified two weeks post-injection.

### *In Vivo* live imaging of CSF dynamics

Two weeks after CLO or Vehicle delivery, macro-scale CSF tracer clearance was evaluated. Mice were anesthetized and prepared stereotaxically as described above. A second burr hole was drilled adjacent to the initial injection site over the right lateral ventricle. A far-red fluorescent tracer, Bovine Serum Albumin conjugated to Alexa Fluor 647 (BSA-647; 2 mg/mL, Invitrogen), was infused into the ventricle (8.5 μL total volume at a rate of 0.5 μL/min).

Immediately following the start of the tracer infusion, the mice were positioned under a specialized fluorescent macro-imaging system equipped with a back-illuminated EMCCD camera. In vivo imaging was then performed using a Zeiss AxioZoom V16 stereomicroscope to monitor the distribution and dynamics of the BSA-647 fluorescent tracer across the brain. Images were acquired every 10 minutes on the Cy5 channel over a total period of 60 minutes (at 20, 30, 40, 50, and 60 minutes post-injection). Image analysis was conducted using ImageJ software to quantify both the percentage of BSA-647– positive area and the total fluorescence intensity throughout the brain. The percentage of tracer-positive surface area and total integrated fluorescence intensity across both dorsal and ventral views were calculated at the 60-minute mark using predefined threshold tracking constraints within ImageJ (NIH). Following imaging, mice were euthanized by decapitation, and brains were collected and fixed overnight in 4% paraformaldehyde (PFA).

### Tissue Processing and Preparation

Exactly 60 minutes after the i.c.v. injection of the BSA-647 tracer, mice were euthanized by decapitation, and brains were collected and fixed overnight in 4% paraformaldehyde (PFA). For mice used in Figure 1, transcardial perfusion was performed to preserve tissue architecture. Briefly, animals were deeply anesthetized and perfused with 20 mL of ice-cold 0.01M phosphate-buffered saline (PBS), followed by 40 mL of ice-cold 4% paraformaldehyde (PFA) in 0.1M phosphate buffer (pH 7.4). Brains were carefully removed from the skull, ensuring the meningeal layers remained intact. Brains were post-fixed in 4% PFA at 4°C for 24 hours, then transferred to a 30% sucrose solution in PBS for 48 hours for cryoprotection. Tissues were embedded in Tissue-Plus optimal cutting temperature (OCT) compound (Thermo Fisher Scientific), frozen in liquid nitrogen-cooled isopentane, and stored at -80°C. Coronal sections (thickness: 30 μm) through the striatum and cerebral cortex were cut using a cryostat and stored in 24-well plates containing antifreeze solution (40% 1× PBS, 30% glycerol, 30% ethylene glycol) at −20 °C until further immunofluorescence use.

### Immunohistochemistry and Quantitative Confocal Microscopy

For immunofluorescence staining, three sections per mouse were processed. Sections were first washed in PBS three times (10 min each) and then permeabilized in 0.5% PBST (1× PBS with Triton X-100) for 20 min, followed by three additional PBS washes (10 min each). Sections were subsequently blocked in 5% goat serum in 0.1% PBST for 1 h at room temperature and incubated overnight at 4 °C with the following primary antibodies: CD206 (rat monoclonal, 1:100, MCA2235; Bio-Rad), Lyve1 (rat monoclonal, 1:100, ALY7; eBioscience), EM48 (mouse monoclonal, 1:200, MAB5374; Sigma-Aldrich), Iba1 (rat monoclonal, 1:500, ab283346; Abcam), and GFAP (rabbit monoclonal, 1:500, 80788; Cell Signaling Technology).

The following day, sections were washed three times in PBS (10 min each) and incubated with appropriate fluorophore-conjugated secondary antibodies (Invitrogen) for 2 h at room temperature. After three additional PBS washes (10 min each), sections were mounted using Fluoromount-G mounting medium containing DAPI.

High-resolution z-stack images were captured using a laser scanning confocal microscope. For BAMs quantification, the linear density of leptomeningeal BAMs was calculated along the outer meningeal borders and expressed as cells per millimeter (cells/mm). Perivascular BAM numbers were quantified within deep parenchymal vessels and expressed as cells per square millimeter (cells/mm²). For mHTT aggregate analysis, nuclear versus extranuclear inclusions within the striatum were differentiated based on spatial co-localization with DAPI boundaries using an automated particle counting macro in ImageJ. For gliosis quantification, the percentage of positive fluorescent area, total integrated fluorescence intensity, and absolute counts of Iba1-positive microglia and GFAP-positive astrocytes were evaluated across three non-overlapping fields of view within the striatum and cortex per animal.

### Statistical analysis

Individual data points are shown in all figures. Investigators were blinded to sex and genotype during data analysis. Statistical analyses were performed using Prism software (version 10.2.3; GraphPad Software, Boston, MA). Data were analyzed using two-tailed unpaired t-tests or two-way ANOVA followed by Sidak’s or Turkey’s multiple comparisons test as indicated in the figure legends. A p-value < 0.05 was considered statistically significant. Significance levels are denoted as follows: ns, not significant, p < 0.05, **p *< 0.01, **p < 0.001, and ***p < 0.0001.

## Author contributions

Y.J: Conceptualization, experimental design, data collection, methodology establishment, statistical analysis, generating figures, and writing original draft. R.S and H.M: Histology, data collection and analysis. E.M: Data curation, data analysis. W.D: Conceptualization, supervision, funding acquisition, project administration, writing, review, and editing manuscript.

## Declaration of generative AI use

During the preparation of this manuscript, the authors used ChatGPT for language refinement and editorial assistance. After using this tool, the authors reviewed and edited the content as needed and take full responsibility for the accuracy and integrity of the manuscript. Generative AI tools were not used to generate experimental data, perform data analysis, or draw scientific conclusions.

## Figure Legend

**Supplementary Figure 1.**
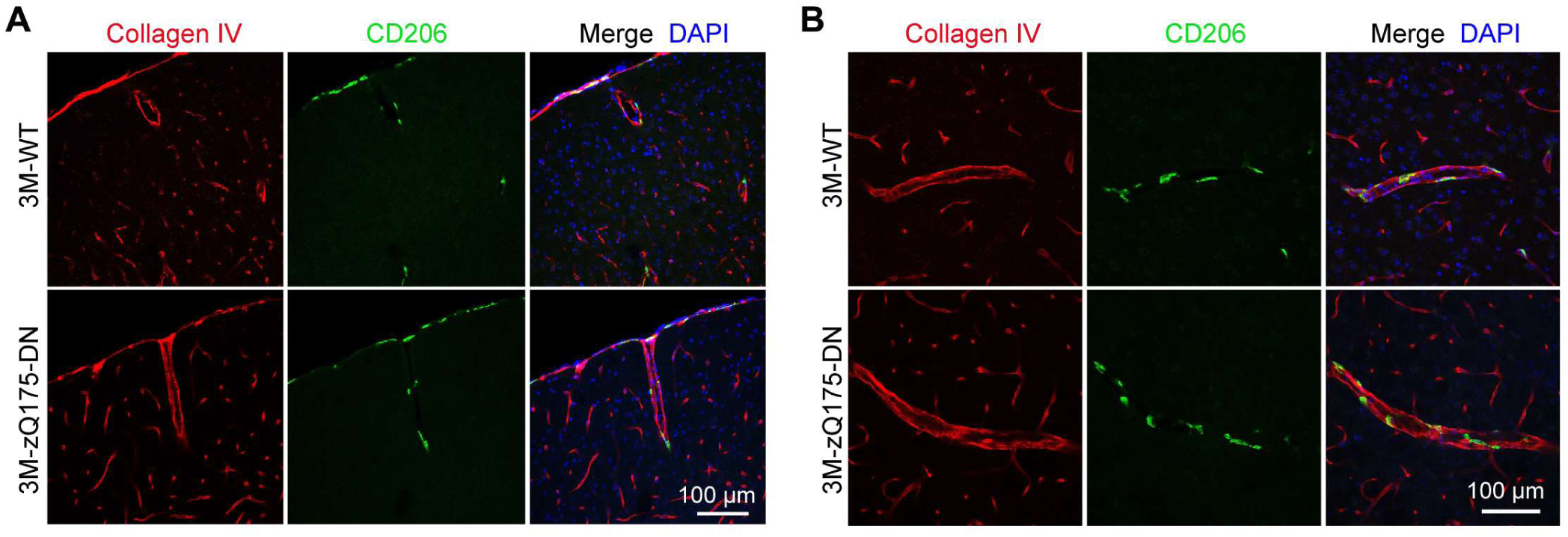
Localization of CD206⁺ BAMs in the brains of WT and zQ175DN mice. **(A)** Representative Collagen IV and CD206 co-immunostained images showing BAMs located in the leptomeninges and along blood vessels in the brains of 3-month-old WT and zQ175DN mice. **(B)** Representative Collagen IV and CD206 co-immunostained images showing BAMs along blood vessels in the brains of 3-month-old WT and zQ175DN mice.

